# Online, Interactive Tutorials Provide Effective Instruction in Science Process Skills

**DOI:** 10.1101/153015

**Authors:** Maxwell Kramer, Dalay Olson, J.D. Walker

**Author notes:** **Corresponding author:** Maxwell Kramer; 5-220 Moos Tower, 515 Delaware Street SE, Minneapolis, MN 55455; (; (department. (MK and DO contributed equally).

## Abstract

Explicit emphasis on teaching science process skills leads to both gains in the skills themselves and, strikingly, deeper understanding of content. Here, we created and tested a series of online, interactive tutorials with the goal of helping undergraduate students develop science process skills. We designed the tutorials in accordance with evidence-based multimedia design principles and student feedback from usability testing. We then tested the efficacy of the tutorials in an introductory undergraduate biology class. Based on a multivariate ordinary least squares regression model, students that received the tutorials are predicted to score 0.824 points higher on a 15 point science process skill assessment than their peers that received traditional textbook instruction on the same topic. This moderate but significant impact indicates that well designed online tutorials can be more effective than traditional ways of teaching science process skills to undergraduate students. We also found trends that suggest the tutorials are especially effective for non-native English speaking students. However, due to a limited sample size, we were unable to confirm that these trends occurred due to more than just variation in the sampled student group.

## Introduction

### Science process skills are an important component of undergraduate biology curricula

A primary goal of undergraduate biology education is to have students develop the ability to think like a scientist. That is, students must develop the ability to ask and answer meaningful biological questions, the core of scientific inquiry. To achieve this goal, students need to master an underlying set of skills including the ability to ask a testable question, propose a hypothesis, design an experiment, analyze data, draw conclusions from evidence and communicate findings. We refer to these skills as *science process skills.* Recent reports by the American Association for the Advancement of Science [AAAS] (2009) and other biology education leaders (Wright & Klymkowsky, 2005; Krajcik & Sutherland, 2010) emphasize teaching these skills as a key goal in improving undergraduate education. Faculty also overwhelmingly value these skills in their students, but traditionally neglect to include them in course design due to a fear of losing time devoted to important subject content (Coil *et al.,* 2010). Here, we offer an alternative approach to incorporating science process skills into course curriculum. We created a series of interactive online tutorials that explicitly teach science process skills and supplement classroom learning with minimal added effort from instructors.

Recent attempts at re-designing undergraduate biology courses show that placing an explicit emphasis on science process skills leads to both gains in the skills themselves and, strikingly, greater retention of subject content. Students who participated in a supplemental course at the University of Washington that taught science process skills and other aspects of scientific culture earned higher grades in introductory biology classes than their peers who did not participate in the course (Dirks & Cunningham, 2006; Buchwitz *et al.,* 2012). Further supporting the benefits of learning science process skills, students’ scores on a science process skills assessment taken after the supplemental course correlated with their introductory biology course grades. The connection between science process skills and overall course grade underscores the impact of explicit instruction in science process skills. In another example, students at Brigham Young University participated in a re-designed cell biology course that placed explicit emphasis on data analysis, interpretation and communication. These students showed improvement in both data analysis and conceptual problems during the course (Kitchen *et al.,* 2003). They also scored higher than students in the traditional content-focused cell biology course on both analysis and recall problems. Other instructors have focused on using primary literature to emphasize critical thinking and science process skills in redesigned courses. The CREATE method, developed at the City University of New York, teaches the nature of science through a series of primary research papers produced by a single research group covering a specific topic (Hoskins *et al*, 2007). Students use active learning approaches to apply science process skills like hypothesis generation, experimental design, data analysis and scientific communication. This approach has yielded gains in critical thinking, gains in experimental design, and improved student attitudes about science in a broad range of postsecondary settings, from community colleges to graduate programs (Hoskins *et al.,* 2011; Gottesman & Hoskins, 2013; Stevens & Hoskins, 2014; Kenyon *et al.,* 2016). While the well-defined CREATE method has proven highly effective in most implementations studied, some instances show no extra gains compared to other active, literature-focused pedagogies (Segura-Totten & Dalman, 2013) In each of these examples, explicit instruction in science process skills led to greater facility with those skills and better content learning in the subject area. Taken together, these results illustrate a clear value in emphasizing science process skills for undergraduates.

### Barriers to teaching science process skills

While the value of teaching science process skills becomes clearer every year, significant barriers still prevent their incorporation into undergraduate curricula. Chief among the barriers is the time commitment required for instruction. As any instructor knows well, time with students is limited and instructors must allocate class time for maximum student benefit. This leads to the familiar debate between covering as much subject material as possible versus a “more is less” approach that focuses on skill development and inquiry while covering a narrower range of topics. Up to this point, most incorporation of science process skills has focused on course-wide redesigns and long-term instruction, as described above. These large-scale changes require significant investment of the instructor’s time and re-allocation of class time. Other attempts at emphasizing science process skills have focused on their integration into laboratory and research experiences, an option not available to all courses and instructors (DebBurman, 2002; Shi *et al*., 2011; Brownell *et al*., 2015; Woodham *et al.,* 2016). Short of these large changes, could smaller scale incorporation of science process skills instruction prove useful to student learning? Little research has evaluated the effect of limited, stand-alone instruction in science process skills.

### Deficiencies of textbook-based instruction

As often as not, students’ only exposure to science process skills comes as an assigned reading in the first chapter of a science textbook. While noble in their intent, these readings often fail to engage students. Students struggle to synthesize the large amounts of information presented in a textbook into an organized framework. This occurs because the amount of processing required by the textbook reading exceeds the capacity of students’ working memory, the amount of information a person is capable of processing at a given time. In psychology terminology, this problem with textbook-based learning is called cognitive overload (Mayer & Moreno, 2003). Static text and layout require readers to devote a large portion of their cognitive load to aspects of the textbook other than the intended content. It is common to find topics covered in one section referencing diagrams found on different pages, forcing students to flip back-and-forth in order to match the text with its graphical representation. The process of moving from page to page requires that the reader split their attention and use what limited working memory they have to synthesize information found in two distinct areas. Additionally, textbooks often overload students with too much information, making it difficult for them to identify and focus on the information worth remembering. While high cognitive load is a common problem for textbooks, not all textbooks are poorly designed. If designed correctly, textbooks can be as effective at teaching students as alternative multimedia. Unfortunately, when it comes to science process skills, this is more often the exception than the rule.

### Benefits of interactive digital instruction

Many students opt out of reading the textbook and instead rely only on in-class lectures to help them learn important topics covered in the course. How can instructor convince their students of the value of pre-class learning? Resistance to learning outside of the classroom is being met with new ways of engaging students. Online, multimedia-based learning is improving education by engaging students through interactive multisensory learning. Harnessing the power of audio, visuals, text, animation and user interactions, multimedia design capitalizes on a variety of ways to deliver information. Students are able to receive and process information through two primary channels: audio and visual. By simultaneously tapping into both channels students are capable of processing larger amounts of information resulting in increased retention. By using this medium, instructors can effectively reduce cognitive load for their students and enable quicker, better-retained learning (Mayer & Moreno, 2003; Evans & Gibbons, 2007; Mayer, 2008; Domagk *et al.,* 2010). A multitude of examples show that online multimedia learning not only helps students to improve their understanding of the concepts presented, but also allows them to integrate new concepts with their existing knowledge base (Carpi, 2001; Carpi & Mikhailova, 2003; McClean *et al.,* 2005; Silver & Nickel, 2005; Chudler & Bergsman, 2014; Goff *et al.,* 2017).

### Interactive digital modules created to teach science process skills

This paper will outline best practices for creating useful and engaging multimedia based tutorials that provide students with an alternative method of learning key material outside of the classroom. Using these principles, we created a series of seven interactive digital modules, each addressing a different science process skill and incorporated them into in a large enrollment introductory biology course. We then assessed the tutorials’ effect on students’ ability to apply those skills compared to students that received only traditional textbook-based instruction. Although we tailored our design strategy to online, multimedia-based learning, instructors can, and should, be able to apply many of these same principles to lecture based design as well.

## Tutorial Design

### Tutorial Design Methods

To create online tutorials, we used Articulate Storyline 2 software (Articulate Global, Inc.), with VideoScribe software (Sparkol Limited, Bristol UK) to create animated whiteboard style videos. All tutorial modules were administered as SCORM 2004 packages via Moodle learning management system, version 3.0.

### Multimedia Design Principles

Our online, science process tutorial was broken up into seven modules outlining the different steps within the scientific process. Although we understand that science does not always progress linearly, we designed these tutorials to be completed in a sequential order. Following the principle of backwards design (Wiggins & McTighe, 2005), we first established a concrete set of learning objectives for each of the seven modules (Supplemental Material). Each module covered two to five learning objectives guided by Bloom’s Taxonomy principles in order to promote higher-level, critical thinking and analytical skills (Bloom *et al.,* 1956; Crowe *et al.,* 2008).

Once completed, these objectives became the roadmap that guided both the content as well as the assessments. When choosing the format for the tutorial, we felt it important that our students experience the process of scientific discovery firsthand. For many years, educational research has supported the use of stories and case studies to teach and make a personal connection with science students (Martin & Brouwer, 1991). Therefore, our science process tutorial follows the storyline of two Nobel laureates, Dr. Barry Marshall and Dr. Robin Warren and their quest to discover the underlying causes of ulcers. From reading primary literature to establishing experiments to analyzing and presenting data, the goal of the online tutorial was to simulate the process of science through the lens of two experienced researchers. Importantly, our online platform offered students an opportunity to learn through interactive engagement rather than passive reading of text.

Having selected learning objectives and the overarching narrative structure for the tutorial, we next turned to designing how the students would interact with the tutorial. A growing body of evidence suggests that multimedia platforms, if properly designed, are effective tools for learning scientific material. For each of the seven individual modules, we followed evidence-based principles of multimedia design outlined, in large part, by Richard Mayer with special attention placed on the coherency and redundancy principles (Mayer, 2008) (Supplemental Materials).

The coherency principle suggests limiting the use of visuals that do not support the overall learning objectives of the project. Every item occupying the learning space increases cognitive load on the learner. Therefore, to maximize learning gains and limit cognitive overload we carefully selected only diagrams and text that support the learning outcomes of the scene. Close adherence to the coherency principle was especially important for this tutorial because of the limited screen space. For each visual scene presented to students, we were careful to limit written text. This approached especially relied on narrated scenes, so that audible narration was not repeated by verbatim text, a concept that closely aligns with the redundancy principle.

The redundancy principle strives to limit the use of printed text for a *narrated* graphic. In other words, narration should describe an image rather than re-reading words on a screen. Simultaneous presentation of words and narration can overload the user’s working memory, making it harder to learn the topic at hand. Instead, research suggests that it is better to replace the words with an image that displays the narrated process.

Finally, to reduce cognitive load, we created a consistent user experience throughout the tutorial (Mayer, 2008; Blummer & Kritskaya, 2009). A consistent navigation interface with clearly labelled buttons allowed student to focus more of their mental capacity on the content presented. We also included repeated prompts and signals that communicate to students what information is most important, when a section is complete, what resources are available and when they are required to make a decision. Approaches as simple as highlighting key words and using a consistent “NEXT” button allow students to reduce extraneous processing.

### Tutorial Audience

We designed our tutorials for undergraduate students with little or no background in the biological sciences. Target populations included students majoring in biology, other science majors and students majoring in non-science subjects. Because the tutorials address fundamental science process skills through the lens of biology, they are appropriate for this broad audience. We also were interested in designing the tutorials to aid students that are non-native English speaking (NNES) students. Interactive and online learning methods can be especially useful for NNES students because they allow students to control the pace of their learning, immediately repeat difficult material, and use visual representations that do not depend on written text. Simulations and learning games are especially effective for NNES students (Abdel, 2002). As such, we recognized the potential our tutorials hold for NNES students and emphasized these elements in our tutorials to make them maximally useful.

### Tutorial Format

In agreement with these design principles, each module followed a similar layout and students quickly became familiar with the general types of slides: the introductory slide, the challenge questions and the summary slides. The introductory slide was the first slide in each module (Figure 1A). On each introductory slide, students watched a brief video that summarized previous modules and outlined upcoming learning objectives. In accordance with the redundancy principle, we used whiteboard-style animations to build the introductory videos (Türkay, 2016). In the videos, drawings appeared as the narrator describes them, an approach that has been shown to increase student engagement (Guo *et al.,* 2014) and that allowed the viewer to take in related information through both the auditory and visual channels simultaneously. Capitalizing on the use of both auditory and visual channels limited cognitive overload and freed the learner to process and store larger amounts of information.

**Figure 1:**
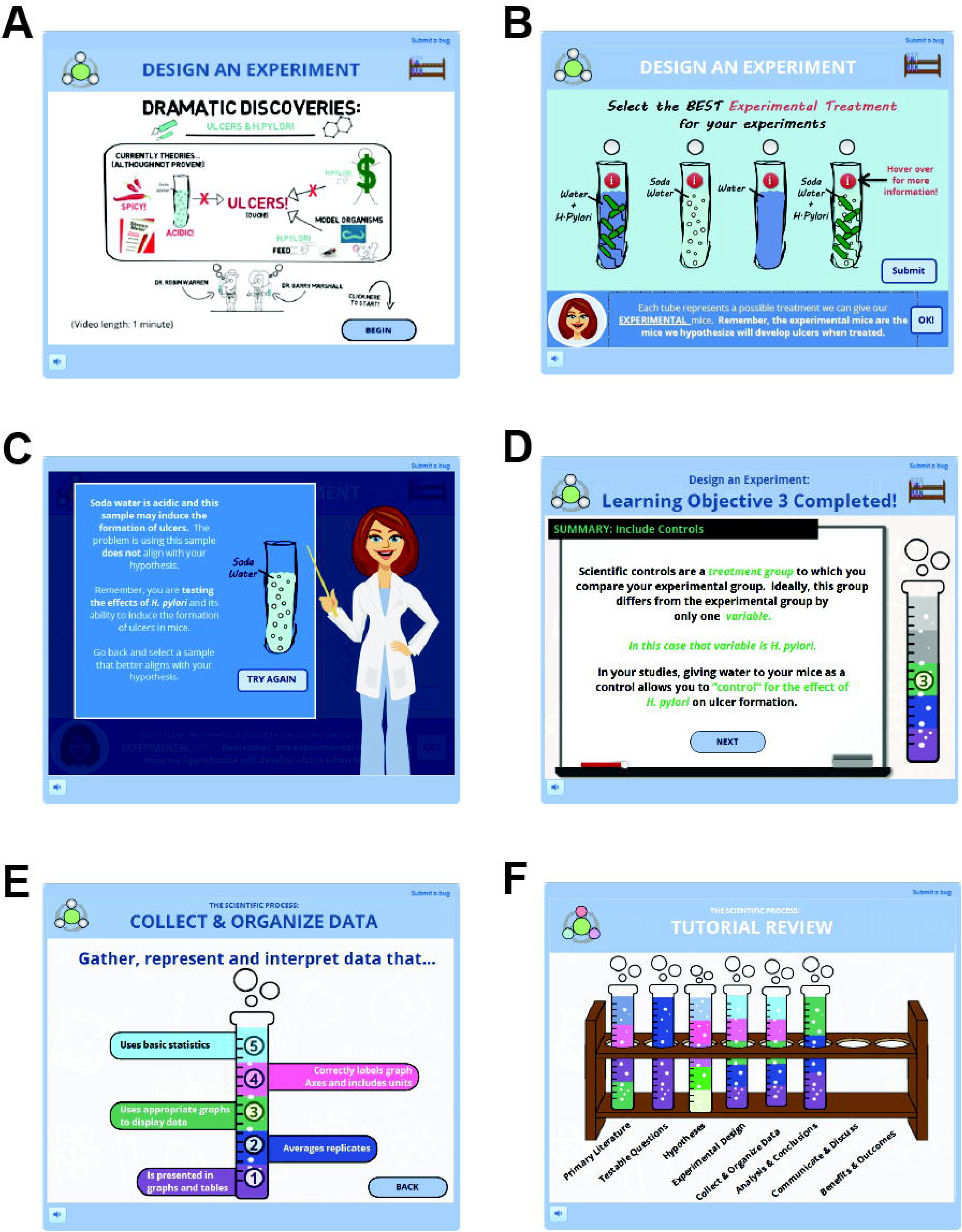
Example screenshots from ‘Experimental Design’ interactive tutorial. Interactive tutorials were designed using consistently formatted interactions. These included module introductions (A), challenge questions (B), and specific feedback (C). Tutorials also include test tube graphics that allow students to track their progress through each module, review individual learning objectives (D), review entire modules (E) and review their progress through the entire tutorial (F). The test tube rack containing test tubes for all completed modules is accessible at any time during the tutorial by clicking on the small rack icon in upper right corner of the tutorial interface. This allows students easy access to review past content and track progress.

Following the introductory whiteboard video, an onscreen character greeted the student and introduced the specific activities for that module. Personalizing interactions between an onscreen character and the learner created a sense teamwork. The onscreen character in each module served to guide the storyline, provided feedback, and prompted the students to think deeply about the learned material (Figure 1B-C). In the module “Design an Experiment,” students were asked to help the researchers develop an experimental outline for the project. As the students progressed through the module, they learned the importance of selecting a model system, assigning proper treatment and control groups, and creating a protocol. Challenge questions posed by the onscreen character during each of these steps pushed students to think critically about typical problems faced at each step of the scientific process.

The tutorials were designed so that when students made choices or answered questions, they reflected on why they chose their specific option and why the other options were better, worse or totally wrong. For example, when selecting the treatment group for their experiments, students chose from four different treatment options. In this scenario, students must select *and* justify the use of their selected treatment. If students selected an incorrect treatment, they received specific feedback that helped guide them to the correct answer (Figure 1C). This feature of the tutorials comports with research indicating that explanatory or informed-tutoring feedback, which provides context and explanations for why an incorrect choice is incorrect and a correct one is correct, is more helpful in the learning process than feedback that is merely corrective (Moreno, 2004; Narciss, 2004).

Lastly, once the student selected the correct answer, we used a consistent format to summarize the question and responses. This interactive slide let students review their correct answer and each of the incorrect answers, including explanations about how each answer fit with the larger concept. In pilot studies, students found the guided feedback very useful for helping them break down complicated material into single units of information. Additionally, we found that asking students to justify their selection forced students to think critically about the question at hand instead of randomly selecting an answer.

Students are better able to engage with multimedia tutorials when they can track their progress and see how close to finishing they are. Some tutorials display the percent complete or the total number of slides. For our tutorial, we created a test tube graphic to represent progress in the tutorial. The test-tube also served a second purpose, to show students the learning objectives for that module. As students complete learning objectives, the test tube fills up (Figure 1D). At the end of the tutorial, the student finishes with a full test tube that displays all of the learning objectives (Figure 1E). This visual tracking method also allows students to “collect” test tubes for all seven modules in the tutorial, creating a game-like incentive that engages students (Figure 1F). Based on student feedback, students appreciated the ability to track their progress through the tutorial.

As mentioned, the test tubes served a second purpose as a way for students to explicitly see the learning objectives for the current module. Upon completing a module, students saw the full test tube with each learning objective filled in (Figure 1E). Each learning objective was as a clickable item that led to a small review activity on that topic. Along with this end of module review, students were always able to review test tubes and learning objectives from previously completed modules by clicking on a consistent icon in the corner of the tutorial screen. This approach requires that students actively seek out necessary information, referred to as “pulling” information. This design strategy was repeated throughout the tutorial as a way to further engage students and give them a sense of control as they work through the tutorial. By using the test tube format, the modules clearly articulate learning objectives and offer a convenient way to revisit previous material.

### Usability testing

After the initial design, we refined our tutorials based on usability testing. Because students experienced the tutorials on their own time and without direct input from the instructor, it was imperative that the tutorials were easy to use, engaging, accessible to students of all abilities, and met our intended learning objectives. To reach these goals, we collaborated with the University of Minnesota Usability Lab. The usability lab provides a physical space to observe a person using a product or design in real time and a process to help improve the effectiveness and accessibility of that product based on those observations. During this process, we met with a usability expert from the lab, determined testing goals, conducted focus groups with undergraduate students, evaluated the results, and decided on specific improvements to the tutorials that addressed identified problems. We chose to focus on improving navigation, optimizing information placement, and determining whether tutorial content was properly challenging for undergraduate students. Students selected for participation were observed as they attempted to complete the “Design an Experiment” tutorial module, immediately followed by a debriefing interview with the usability expert. During the testing session, we directly observed students via a one-way mirror, their computer screen via simulcast, and their eye movements via eye-tracker camera and software. We tested tutorials in this way with six undergraduate students, including both biology majors and non-majors, and native English speakers and non-native English speakers.

During usability testing, we found that students were most engaged by concise delivery of information with emphasis placed on key concepts (*e.g.* bold, italics, or color), extremely consistent visual markers for navigation, and immediate feedback for wrong answers to challenge questions. As expected, the level of difficulty, as measured by in tutorial assessment questions, was higher for students who were majoring in subjects outside of the sciences. Lastly, we found that distinct aspects of the tutorials engaged non-native English speakers. Consistent layout and visual design were important for easy navigation. We suspect that this design allowed students to spend less cognitive energy decoding the instructions and navigation, and to focus more on absorbing content from the tutorials. Along similar lines, non-native English speakers also specifically appreciated repeated presentation of key ideas and optional chances to review those important concepts. We incorporated all of these observations into the design strategy for the final version of the tutorials (see Supplemental Material for specific feedback).

## Research study

### Design

To investigate the effectiveness of our online science process skills tutorials, we conducted a quasi-experimental study in a large format introductory biology course for students majoring outside biology in spring of 2017 at the University of Minnesota, a large public research institution in the Midwest of the United States (Figure 2). Without prior knowledge of this study, students enrolled in one of two sections, each with the same instructor. One section was assigned the tutorials as an out-of-class activity to supplement textbook reading over the first two weeks of the course, the “online tutorial” group (n=118). Student completion of the tutorials was high: 93% of enrolled students completed at least three out of five tutorials and 88% of enrolled students completed all five tutorials. The other section was assigned only out-of-class textbook reading covering similar subject material over the same two weeks, the “textbook reading” group (n=118). For reasons of fairness, students in the textbook reading group were given access to the online tutorials following the experimental period and assessment. Students were incentivized to complete the tutorials with a small amount of homework credit (low stakes). The University of Minnesota Institutional Review Board deemed this study exempt from review (Study Number: 1612E01861).

**Figure 2:**
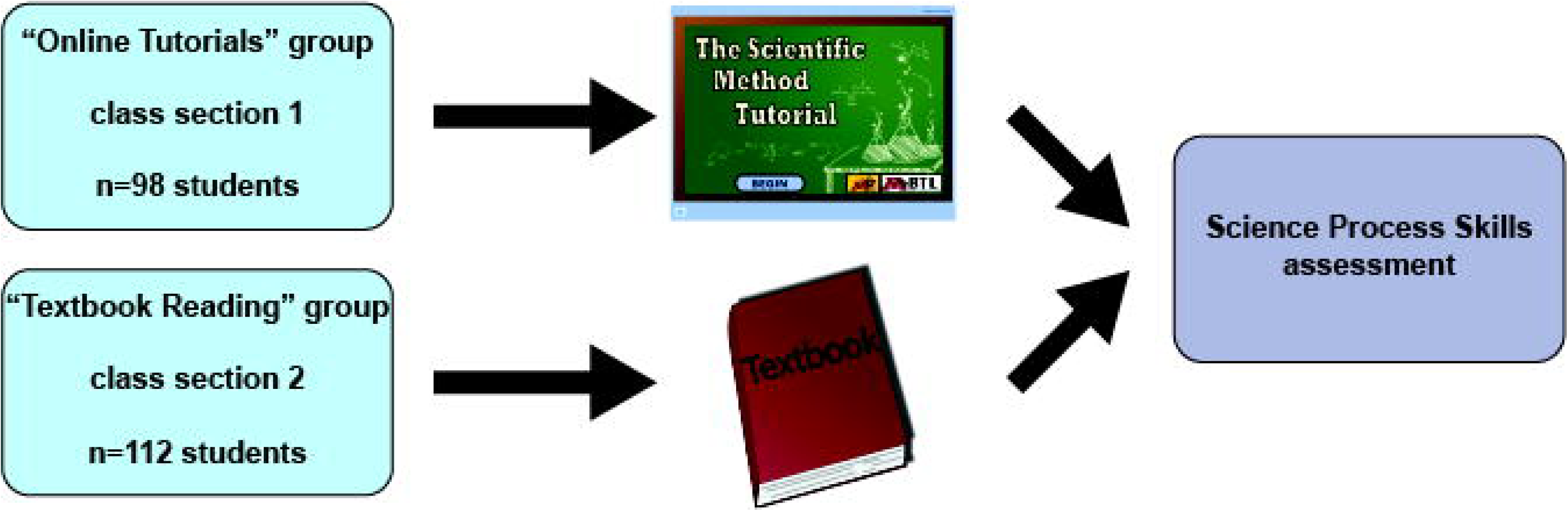
Experimental design. Students enrolled in parallel sections of the same introductory biology course were given two weeks to complete either the online tutorials or textbook reading. At the end of the two weeks, a 15 question quiz assessed students’ science process skills.

### Measures

While recent studies in undergraduate education have focused on ways to improve scientific literacy and proficiency, the ability to test the effectiveness of these interventions is limited. To date, very few formative assessments designed to gauge science process skills have been validated, and those that have, are geared towards assessing K-12 students. Therefore, due to the lack of validated assessments, we chose to modify a well-accepted science process skills assessment tool known as the Integrated Process Skills Test, or TIPS II (Burns *et al.,* 1985). Modified versions of TIPS II have been successfully used to assess science process skills in high school seniors, a population of students very similar to our own (Kazeni, 2005). Rather than design and validate a new assessment tool we chose to select questions from TIPS II for our assessment.

The fifteen-question assessment was designed to assess five different categories of science process skills including graphing and interpreting data, identifying variables, stating hypotheses and selecting operational definitions (Supplemental Material). To populate each category in our assessment we selected only the questions from TIPS II with the highest item discrimination index (as reported in Kazeni, 2005), a statistical measure that distinguishes between high performing and low performing examinees for a given assessment. The average item discrimination index of questions selected from TIPS II was 0.4, well above the acceptable range (>0.3; Bass *et al.,* 2016). Student scores on these fifteen TIPS II items were averaged to create the main outcome variable in this study.

The ability to apply science process skills was assessed in both the online tutorial and textbook reading groups at the same point in the semester, after instruction was complete, using our modified TIPS II test. Student participation in the assessment was high for both groups: 98/118 for the online tutorial group and 112/118 for the textbook reading group.

### Data

The total pool of study participants was 54.3% female and 45.7% male, with a median age of 20.34 years. The respondents were 0.5% American Indian, 8.7% Asian, 6.6% black, 2.9% Hispanic, 15.8% international, and 65.3% white. They were 3.6% first-year students, 40.6% sophomores, 32.0% juniors, and 23.9% seniors, and 18.3% were listed as non-native English speaking (NNES) students. 74.2% of participants were non-science majors.

Because participants were not randomly assigned to the online tutorial and textbook reading groups, it was important to establish comparability of the two groups on the available exogenous variables, which included aptitude variables (composite ACT score, GPA) and demographic variables (ethnicity, sex, age, major, NNES status). We ran appropriate group comparison tests and determined that the two groups of participants were statistically equivalent on all aptitude and demographic variables, with the exception of age: the mean age in the online tutorial group was slightly higher than the mean age in the textbook reading group (20.55 vs 19.88, p = .008).

### Analysis and Results

We constructed a multivariate ordinary least squares regression model to predict participants’ performance on our outcome variable of interest, namely score on the modified TIPS II test. For this model, casewise diagnostics were generated and examined to locate outliers in the data set, defined as cases with standardized residuals greater than 3.3. This procedure revealed three outliers for the dependent variable. On inspection, these cases were not otherwise anomalous, so they were retained in the data set. Variance inflation factor (VIF) statistics were also generated to check for multicollinearity among the predictor variables. In no case was the VIF statistic greater than 1.135, far from the common cut-off of 4, so multicollinearity did not appear to be a problem in the data set. The Durbin-Watson statistic of 2.346 indicated very little auto-collinearity in the data.

The model included just three predictor variables (GPA, NNES status, and treatment group); no other demographic variables were significantly related to TIPS II score. The model was highly significant (p < .001) and accounted for a small-to-moderate amount of the variation in TIPS II score with an R^2^ value of .403, adjusted R^2^ = .140.

The covariate GPA was significantly related at the p < .05 level to TIPS II scores (t = 2.799, p = .006), as was the main treatment of interest, being in the online tutorial group (t = 2.600, p = .010). The predictor variable representing NNES status was negatively associated with TIPS II scores, but it was not statistically significant (t = -1.164, p = .246).

Given the results displayed in Table 1, we can conclude that for each unit increase in a student’s GPA, we can expect more than a 1-point increase in that student’s TIPS II score. Similarly, being in the online tutorial group was associated with a 0.824 point increase in a student’s TIPS II score, while holding other variables in the model constant. The pooled standard deviation for the TIPS II score variable was 1.99, so the effect size for GPA was moderate to large (almost 60% of a standard deviation), while the effect size for being in the online tutorial group was moderate (over 40% of a standard deviation).

**Table 1.**
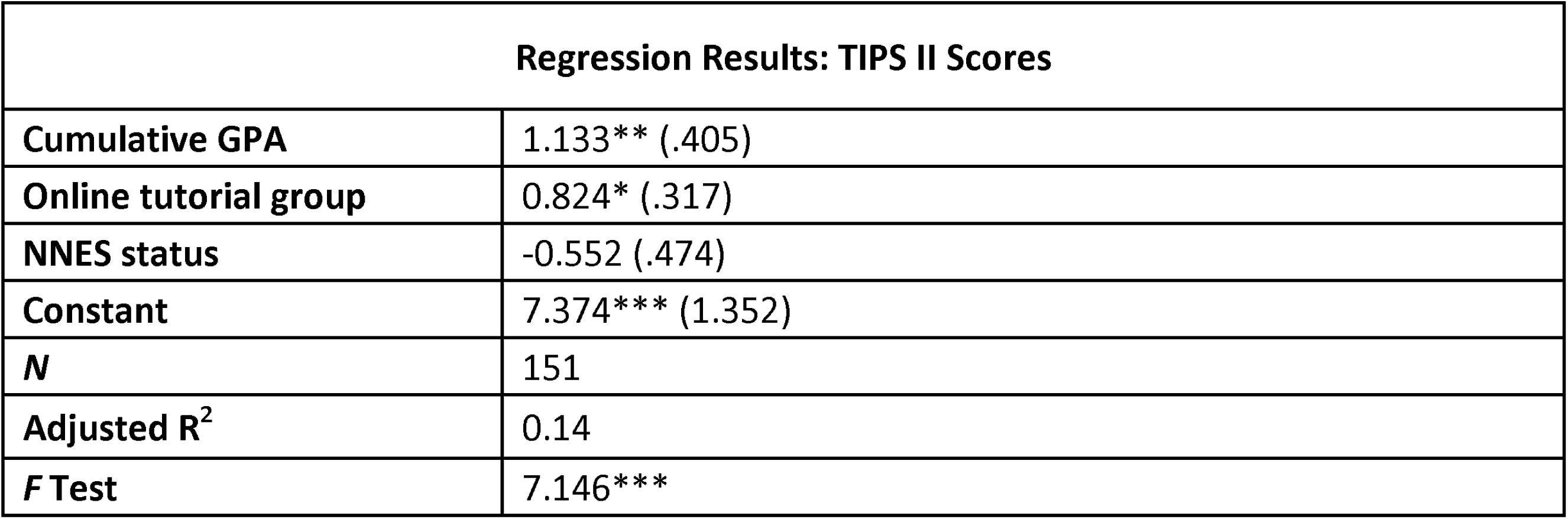
Ordinary least squares regression, with TIPS II score as dependent variable. Cell entries are unstandardized beta coefficients with standard errors in parentheses. *p < .05, **p < .01, ***p < .001

| Regression Results: TIPS II Scores |  |
| --- | --- |
| Cumulative GPA | 1.133** (.405) |
| Online tutorial group | 0.824* (.317) |
| NNES status | -0.552 (.474) |
| Constant | 7.374*** (1.352) |
| *N* | 151 |
| Adjusted R<sup>2</sup> | 0.14 |
| *F* Test | 7.146*** |

### Results for NNES Students

One sub-population of interest in this study was the group of students identified as non-native English speaking (NNES) students, so additional analyses were performed to examine the effects of the online tutorials on this group. There were some indications in the data that the online tutorials may have benefited NNES students to a greater degree than they benefited students who were native speakers of English, but low subgroup N rendered these indications unclear.

First, we asked whether being in the online tutorial group improved the TIPS II scores of NNES students more than it improved the scores of other students. The answer was, nominally, yes (Figure 3). Among native speakers, being in the online tutorial group helped (mean difference 0.731, p = .025). However, among NNES students, being in the experimental group helped *more* (mean difference 1.308, p = .311). The latter difference is over ½ of a standard deviation and is much larger than the difference between the scores of tutorial and textbook native English speakers, but it does not test significant because of low N in the NNES group (20 total).

**Figure 3:**
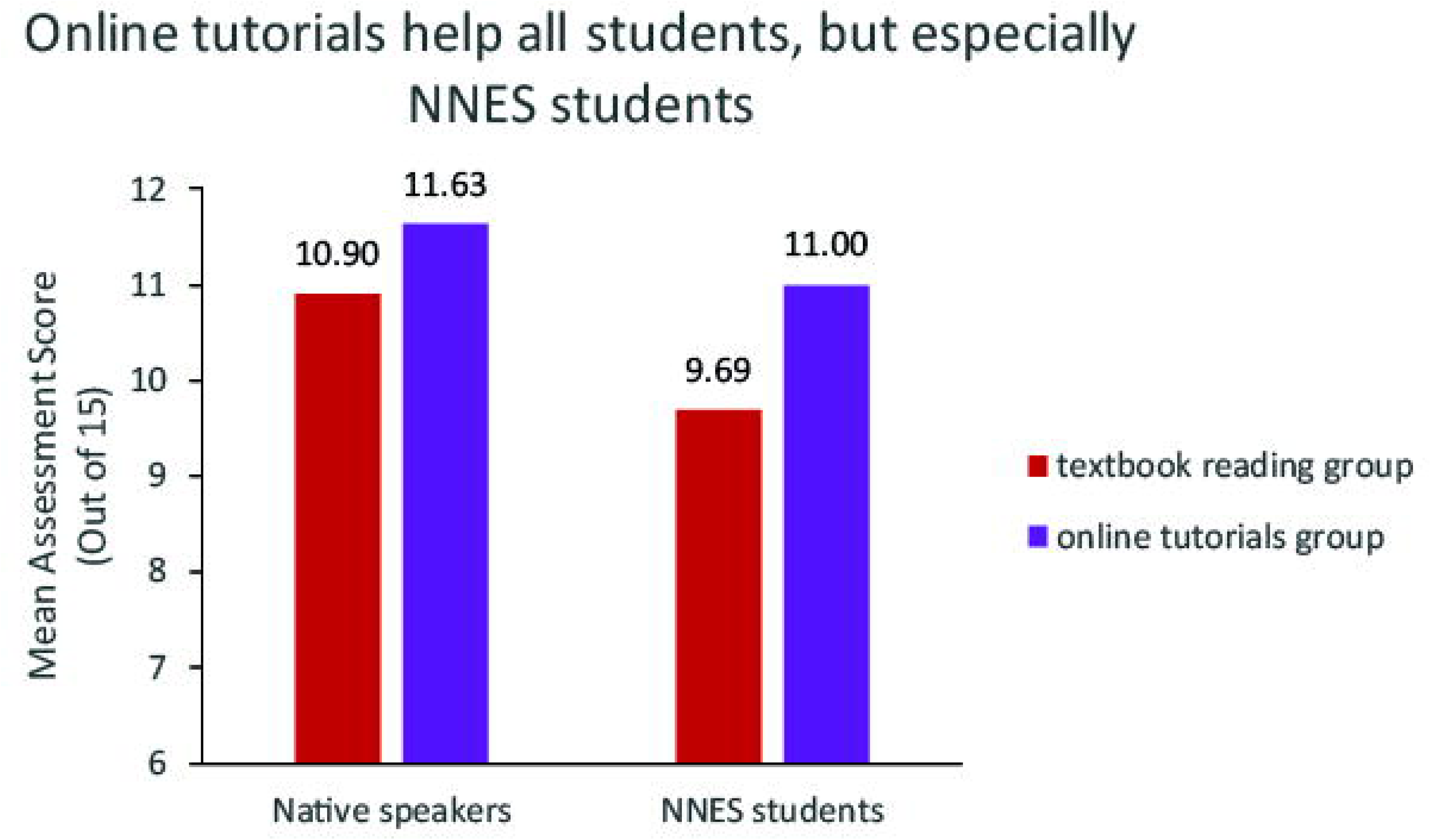
Mean scores for science process skills assessment after textbook (red) or online tutorial (purple) assignment. Comparison highlights difference between instructional approaches for native and non-native English speakers.

Second, we noted that NNES students had average TIPS II scores that were significantly lower than the scores of native English speakers (10.15 vs 11.23, p = .030). So we asked whether being in the online tutorial group helped NNES students to *close the gap* between themselves and native speakers. Again, the answer was, nominally, yes (Figure 4). The difference in average TIPS scores between native speakers and NNES students was 1.21 points in the textbook reading group (p = .101), while in the online tutorial group the difference was just about half of that, 0.633 points (p = .255).

**Figure 4:**
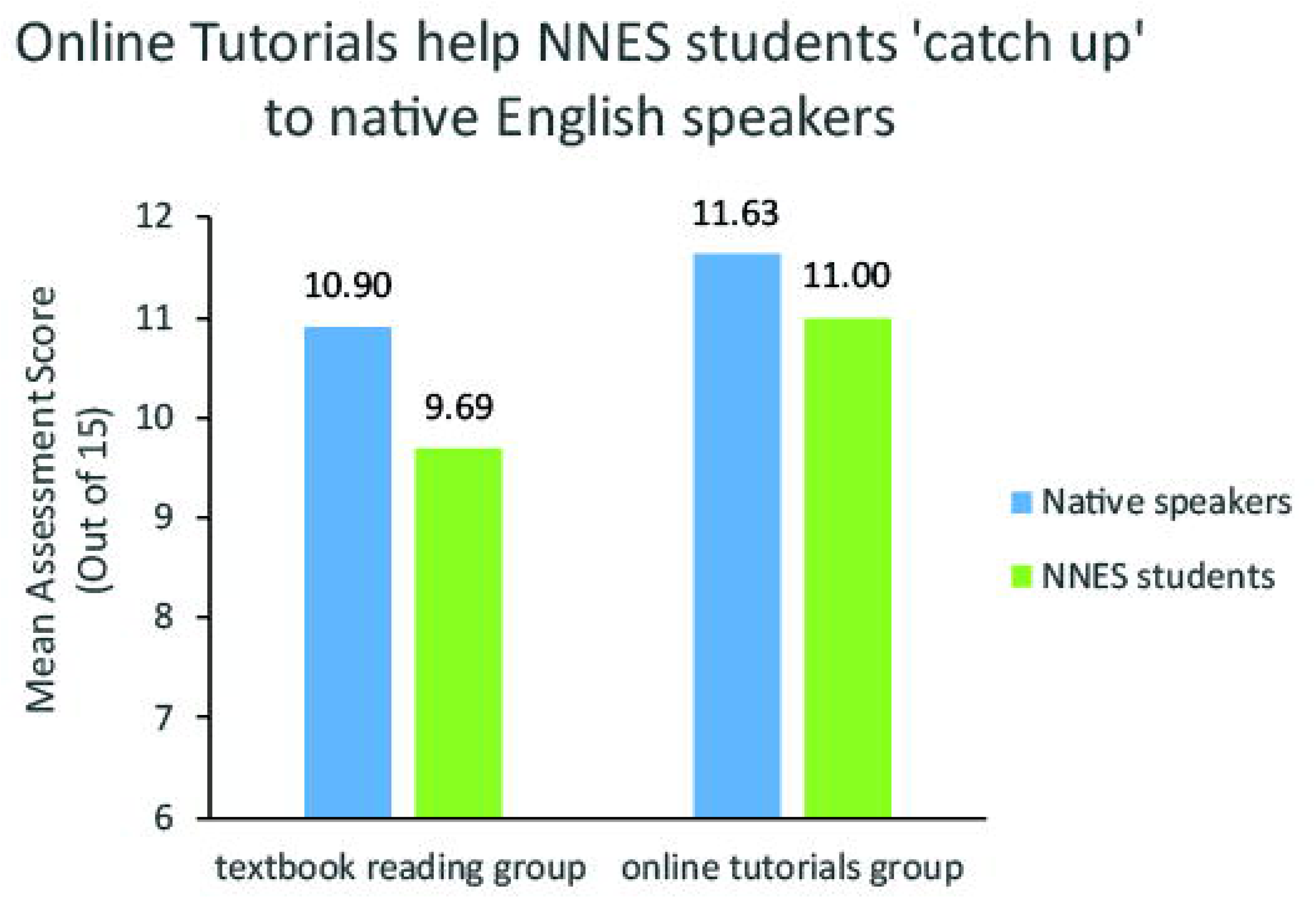
Mean scores for science process skills assessment among native and non-native English speakers. Comparison highlights the difference between the two groups when given textbook reading or online tutorials.

## Discussion

We created a series of online, interactive tutorials with the goal of helping undergraduate students develop science process skills. We designed the tutorials in accordance with evidence-based multimedia design principles and student feedback from usability testing. Based on a multivariate ordinary least squares regression model, students who received the tutorials are predicted to score 0.824 points higher on a 15-point science process skill assessment than their peers that received traditional textbook instruction on the same topic. This moderate but significant impact indicates that well designed online tutorials can be more effective than traditional ways of teaching science process skills to undergraduate students. We also found trends that suggest the tutorials are especially effective for NNES students. However, due to a limited sample size, we were unable to confirm that these trends occurred due more than just variation in the sampled student group.

### Strategies for student engagement

A preponderance of evidence shows that active learning improves student learning compared to traditional lecture approaches (Freeman *et al.,* 2014). Here, we show that even on a very limited scale, a switch from traditional textbook reading to include a more active approach results in improved learning. To achieve this, we designed the tutorials to maximize interaction and engage students in multiple ways in our tutorials. First, the nature of the multimedia-driven, interactive format increases interactivity compared to textbook reading. While completing the tutorial, students are required to make decisions and those decisions shape how the rest of the tutorial proceeds. Multiple sensory channels are engaged in complimentary ways when students see images, hear narration and use computer interfaces to interact with the virtual environment. Each of these features creates opportunities for student engagement. While not every aspect might engage a particular student, the multiple levels of engagement increase the likelihood that one of those elements will grab and keep a student’s attention during the tutorial.

We also chose to use a continued narrative across the modules in the tutorial to create a human connection with students. Our storyline focused on the connection between stomach ulcers and the bacteria *H. pylori* and the story of how two scientists, Robin Warren and Barry Marshall made that discovery in the 1980s. We used familiar settings, like a doctor’s office and library, specific characters, and a medical mystery to create personal connections with students. We suspect that by connecting the scientific method to a human story, students were able to engage more quickly and deeply with the presented skills. While students appreciated the story, it was unclear whether they preferred a more cartoonish or realistic presentation style. Casual feedback from students covered all perspectives from appreciation to distaste for our animation-based style, but NNES students clearly favored the cartoon style. This is consistent with other research that shows schematics are more effective at teaching biological mechanism than realistic images (Scheiter *et al.,* 2009). Comparing different levels of realism would make an area excellent future investigation.

Lastly, we maximized student engagement by offering frequent, specific feedback and opportunity for reflection. We offered feedback in real time as students made choices, after challenge questions and at the end of each module. We found that students appreciated a consistent format for feedback and were able to focus on their thought process and reasoning when presented with a standardized feedback interaction. We also found that feedback needed to be specific. During usability testing, we observed that when the tutorial provided targeted feedback explaining why each of wrong answers were wrong, students stayed more engaged and were more comfortable with the presented concepts at the end of the tutorial. Lastly, we gave students a chance to revisit and modify their ideas. This was especially evident in the “Propose a Hypothesis” module where the tutorial guides students to revise their hypothesis to make it more scientifically sound. These types of interactions provide both engagement and opportunity for formative assessment by the student.

### Potential Uses for Tutorials

In this study, we used our online tutorials as an out-of-class assignment, but they could be employed in a variety of situations. We also created the tutorials with the intention that both the narrative content and the learning goals associated with science process skills could be adapted to suit individual instructor’s needs. In fact, we tweaked several learning objectives before our study to better align the tutorial with the instructor’s other activities and materials. This included modifying language and omitting one previously produced module entirely. We see this flexibility as a strength of the tutorial format. Unlike textbook and other static material, the instructor is free to modify these tutorials to meet their needs.

### Impact on broader student learning and NNES students

One major strength of explicit science process skill instruction is the enhanced student learning in other facets of undergraduate biology education, including content retention (Kitchen *et al.,* 2003; Dirks & Cunningham, 2006; Ward *et al*, 2014). Unfortunately, in this study, we were not able to reliably assess the tutorial’s impact on student learning other than specific gains in science process skills. We look forward to extending our assessment to other facets of student learning. Similarly, based on our design decisions, student feedback during usability testing, and assessment trends, we suspect that the tutorials are especially helpful for NNES students. Better support for this idea, would require a more targeted intervention involving larger numbers of NNES students. As modern biology deepens its international connections and international students continue to enroll at English speaking universities, we hope that the online tutorial format will provide a strong instructional tool to help minimize gaps between native English speaking and NNES students.

## Supplemental Material

Supplement 1: Principles of Multimedia design

Supplement 2: Tutorial Learning Objectives

Supplement 3: Usability testing results

Supplement 4: Assessment questions

Supplement 5: Descriptive statistics for individual assessment questions

## Accessing the Tutorials

All seven of out tutorial modules can be accessed in HTML5 format at (https://sites.google.com/umn.edu/btltutorials). For LMS-compatible files or to discuss modifying modules for your use, please contact the authors.

## Acknowledgments

Dalay Olson and Max Kramer were supported by the University of Minnesota International Student Academic Services Fee and the Howard Hughes Medical Institute. We would like to thank Deena Wassenberg, Robin Wright, Annika Moe, Sadie Hebert, Jessamina Blum and Mark Decker for their input and support. We would also like to acknowledge the Understanding Science website (University of California Museum of Paleontology, <u>http://undsci.berkeley.edu</u>) for their science process skill diagram.

